# Antigen-reactive CD4+ T cells after SARS-CoV-2 vaccination show divergent phenotypic states with or without restimulation bias

**DOI:** 10.1101/2025.11.19.688845

**Authors:** Katharina Kocher, Felix Drost, Christine Schülein, Bernd Spriewald, Benjamin Schubert, Kilian Schober

**Affiliations:** Mikrobiologisches Institut – Klinische Mikrobiologie, Immunologie und Hygiene, Universitätsklinikum Erlangen und Friedrich-Alexander-Universität (FAU) Erlangen-Nürnberg, Erlangen, Germany; Institute of Computational Biology, Helmholtz Zentrum München — German Research Center for Environmental Health, Neuherberg, Germany; School of Life Sciences Weihenstephan, Technical University of Munich, Munich, Germany; Department of Internal Medicine 5, Universitätsklinikum Erlangen and Friedrich-Alexander-Universität (FAU) Erlangen-Nürnberg; FAU Profile Center Immunomedicine, FAU Erlangen-Nürnberg, Erlangen, Germany

## Abstract

Understanding antigen reactivity is crucial for characterizing CD4+ T helper (Th) cell fate, yet conventional peptide restimulation assays introduce phenotypic bias by activating cells *ex vivo*. However, by performing single-cell RNA and T cell receptor (TCR) sequencing on both antigen-stimulated and unstimulated samples, clonotypes can be tracked across conditions to identify antigen-reactive CD4+ T cells and simultaneously be assessed for their phenotypes in the unperturbed state. Using this ‘reverse phenotyping’ strategy, complemented by DNA-barcoded peptide-HLA (pHLA) class II multimers, we here tracked SARS-CoV-2 spike-reactive CD4+ T cells longitudinally after repeated mRNA vaccination. Without stimulation, reactive clones showed more Th-neutral features and less of an activated Th1-like state than would be assessed after antigen restimulation. Furthermore, transgenic TCR re-expression guided separation of antigen-specific from bystander-activated clones. These results uncover unbiased phenotypes of antigen-reactive CD4+ T cells and highlight that cell state classification can differ fundamentally when judged by phenotype versus function.

## Introduction

The fate of a T cell is determined by the specific antigen it recognizes. Accurately identifying antigen-reactive T cells is crucial for understanding immune responses, yet most available methods rely on *ex-vivo* stimulation followed by detection of activation markers or cytokines^1^. While these assays are effective for detecting functional reactivity, they can alter the cells’ native phenotype^2^. and may inadvertently include cells that are activated independently of antigen specificity, known as bystander-activated cells^3^. Therefore, there is a pressing need for approaches that can identify antigen-reactive T cells without introducing such stimulation-induced phenotypic bias. This challenge is particularly pronounced for CD4+ T cells, as generating peptide-human leukocyte antigen (pHLA) multimers for HLA class II is more complex than for class I.

Nevertheless, for pathogens like SARS-CoV-2, many immunodominant HLA class II-restricted epitopes have been characterized^4–11^, such as DPB1*04:01/S_167_, DRB1*15:01/S_751_, S_816_ (potentially presented by multiple alleles) and DRB1*15:01/S870. For some of these epitopes, pHLA class II multimers are available and have been instrumental in characterizing the phenotype of SARS-CoV-2 epitope-specific CD4+ T cells, especially in settings with limited cell numbers *ex vivo*^8,12^. However, pHLA class II multimers are inherently restricted to the analysis of individual epitopes, limiting their ability to provide a comprehensive view of antigen reactivity that broader antigen restimulation approaches can offer. To address these limitations, we previously demonstrated that it is possible to combine the strengths of both strategies by tracking how a T cell clone responds to antigenic stimulation and retrospectively examining its phenotype in the unstimulated state^13^. We termed this method “reverse phenotyping”. Our initial proof-of-concept study, however, was constrained by non-standardized samples collected at varying time points during or after severe COVID-19 infection^13^. Despite these constraints, we corroborated earlier findings of a cytotoxic Th1 (type 1 helper T cell) lineage-driven response of CD4+ T cells to SARS-CoV-2 infection^14–18^.

Importantly, reverse phenotyping revealed that transcriptional signatures of ‘Th1-ness’ (as indicated by expression of the hallmark transcription factor *TBX21*) are overestimated following *ex-vivo* peptide stimulation (or, in fact, underestimated without stimulation)^13^. While this does not negate the Th1 and cTfh predominance in blood – since reverse phenotyping confirmed upregulation of *IFNG* after stimulation, consistent with a functional Th1 definition^19^ – these observations suggest that phenotypic and functional classifications of cell states may yield divergent conclusions. This is particularly relevant given the uncertainty about how closely *ex-vivo* stimulation conditions mimic antigen encounter *in vivo*.

Unlike exploratory snapshot analyses performed after infection, a vaccination setting offers a unique opportunity for standardized, longitudinal investigation of antigen-reactive T cell phenotypes. The widespread SARS-CoV-2 vaccination campaign not only provided significant clinical benefits but also enabled rare, systematic studies of fundamental human T cell responses. Multiple studies have shown that repeated mRNA vaccination induces spike-reactive CD4+ T cells with Th1/cTfh lineage characteristics and central/effector memory (TCM/TEM) differentiation states^9,12,20–25^, mirroring responses seen in SARS-CoV-2 infection. However, these studies were either limited to individual HLA class II-restricted epitopes, or analyzed CD4+ T cell phenotypes after antigenic restimulation *ex vivo*.

Here, we employed reverse phenotyping – complemented by DNA-barcoded pHLA class II dextramers for DPB1*04:01/S_167_ – to longitudinally characterize the phenotype of SARS-CoV-2 spike-reactive T cells both with and without *ex vivo* restimulation. Further validation using TCR-transgenic T cells revealed that, in the absence of restimulation, spike-reactive T cells during the memory phase predominantly exhibit a transcriptional profile consistent with a Th-neutral effector memory (TEM) state, lacking strong Th1 polarization. These Th-neutral signatures are otherwise obscured by the robust induction of Th1 programs observed in assays involving *ex-vivo* restimulation.

## Results

### Study design and analysis approach

Our previously published CoVa-Adapt cohort^26^ consists of 29 healthy donors who received three SARS-CoV-2 mRNA (Comirnaty) vaccinations in a highly synchronized manner (**Fig. 1A; Table S1**). A primary (P) vaccination was given at day 0, followed by a secondary (S) vaccination at day 21 and a tertiary (T) vaccination at day 240. Blood samples were obtained 10 days after primary, secondary, and tertiary immunization (P10, S10, and T10), and additional memory time points were studied at S210, T108, and T189. 12-20 donors were subjected to conventional assessment of spike antigen reactivity through an IFNγ ELISpot. In addition, two donors were analyzed by reverse phenotyping, exposing peripheral blood mononuclear cells (PBMCs) either to overlapping 15mer peptide mixes of the spike antigen *ex vivo* or leaving them unstimulated. Subsequent flow cytometric sorting of non-naïve CD4+ and CD8+ T cells was followed by single-cell RNA sequencing (scRNAseq), including paired TCR sequencing (VDJseq) (**Suppl. Fig. 1A; Table S2**). For selected donors, we also performed epitope-specific scRNAseq analysis of CD4+ T cells with pHLA class II dextramers for the DPB1*04:01/S_167_ spike epitope and CITEseq assessment of 130 surface proteins (**Fig. 1A; Table S3**).

**Fig. 1:**
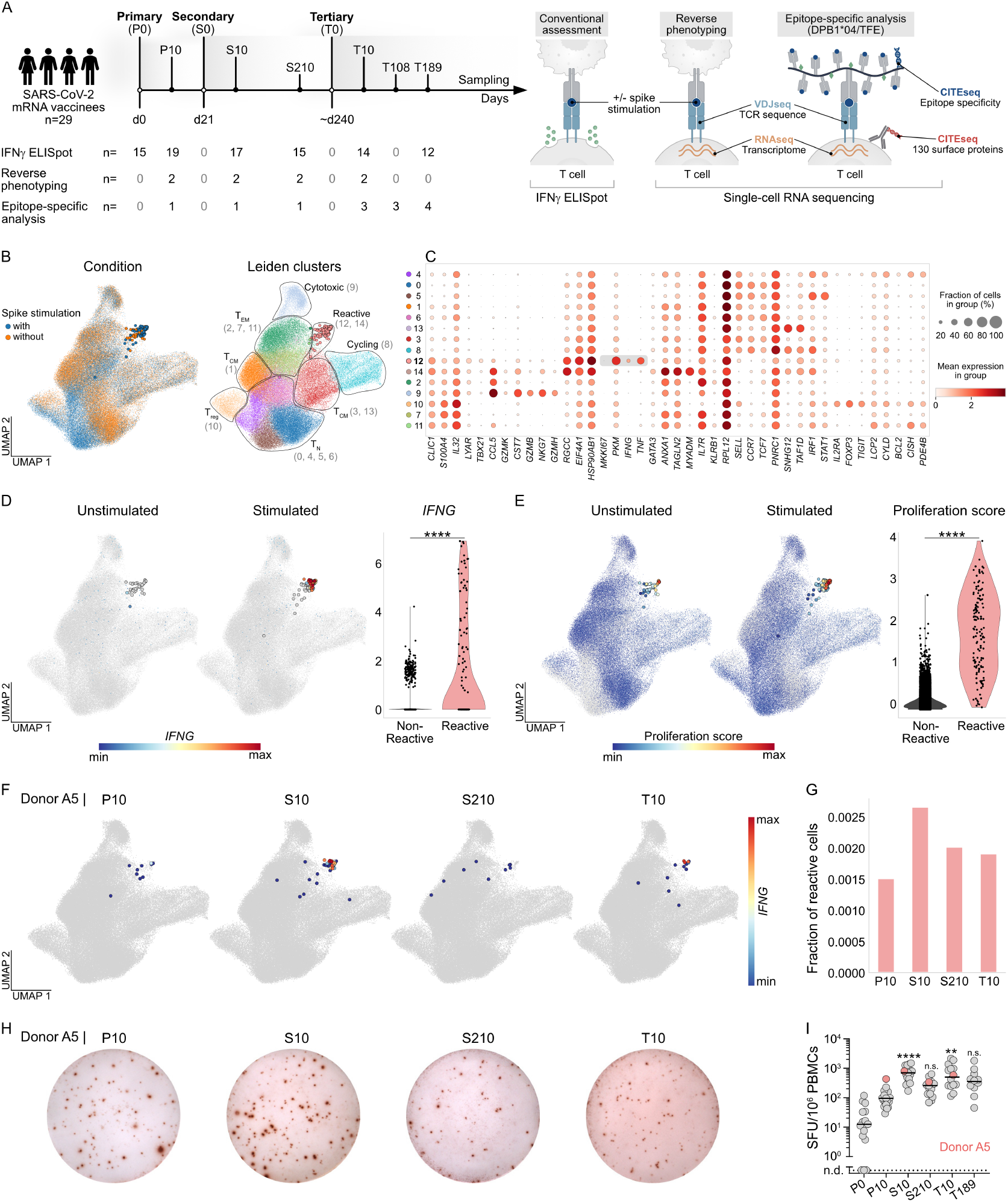
Identification of SARS-CoV-2-specific T cell responses by reverse phenotyping. **A** CoVa-Adapt study design and sample collection scheme. For all donors, PBMCs were collected at day 0 (P0), 10 days after primary (P10), 10 and 210 days after secondary (S10, S210), and 10 and 189 days after tertiary (T10, T189) vaccination. For selected donors, PBMCs were additionally sampled 108 days after tertiary vaccination (T108, n=7). Vaccination-induced T cell responses were characterized for most donors on a quantitative level by IFNγ ELISpot. Selected CoVa-Adapt donors were subjected to in-depth characterization using scRNAseq (reverse phenotyping and epitope-specific analyses) followed by TCR functional testing. **B-G** scRNAseq data from the reverse phenotyping dataset. For reverse phenotyping, PBMCs were re-stimulated with 15-mer peptides covering the complete wildtype spike protein or left untreated. Sorted non-naïve CD4^+^ and/or CD8^+^ T cells were subjected to scRNAseq. Only CD4^+^ T cells are shown (annotation described in methods section). The full dataset is depicted in Suppl. Fig. 1. **B** UMAP of stimulated (blue) and unstimulated (orange) T cells (left panel) and Leiden clusters (right panel; cluster names in UMAP, cluster numbers on the right) (n=101,939 cells in total). Cells located within the reactive cluster are displayed with increased point size. **C** Dot plots of log-normalized expression of representative genes per cluster. Selected genes of the reactive cluster are highlighted in grey. Numbers on the left indicate cluster numbers with reactive cluster 12 highlighted in bold. **D-E** *IFNG* expression (D) and proliferation score (E) in unstimulated (stimulated cells in grey) and stimulated (unstimulated cells in grey) CD4^+^ T cells (left), and quantification in the stimulated condition of cells in the reactive cluster versus all other clusters (right). Cells located within the reactive cluster are displayed with increased point size. For *IFNG*, cells with log-normalized gene expression of 0 are shown in grey in UMAPs. Statistical testing by Mann-Whitney U test. **F** UMAP visualization of cells classified as reactive (cells located in the reactive cluster or belonging to clones where at least one cell is in the reactive cluster) from donor A5 at individual time points after primary (P), secondary (S) and tertiary (T) vaccination in the stimulated condition. Color gradient indicates *IFNG* expression at indicated time points. Non-reactive cells and cells from other donors are shown in grey. **G** Fraction of cells from donor A5 at each time point belonging to the reactive cluster. **H-I** Identification of spike-reactive T cells after 20h of *in vitro* re-stimulation of PBMCs with 15-mer peptides covering the complete wildtype spike protein. Peptides were provided in two subpools, S1 (depicted in Fig. 1H) and S2. Primary data (H) of donor A5 is shown. Quantification (I) of spot-forming units (SFU) for IFNγ ELISpot (combined frequencies of S1 and S2 subpools), data points represent individual donors (n=12-19 per time point), solid lines indicate the mean. Samples without SFU above the negative control were set to not detected (n.d.). Donor A5 is highlighted in pink. Statistical testing by Kruskal-Wallis test followed by Dunn’s multiple comparisons test. Significant differences to P10 time-point are indicated. **p<0.01, ****p<0.0001, n.s. not significant.

### Identification and dynamics of reactive clonotypes

Visualization of single-cell RNA sequencing data and concomitant Leiden clustering revealed one cluster (cluster 27), which was predominantly detected in the stimulated condition and mostly consisted of CD4+ T cells (**Suppl. Fig. 1B-C; Table S4**). This reactive cluster, in contrast to other clusters, showed marked upregulation of activation-related genes such as *IFNG* as well as enhanced expression of proliferation-associated genes (**Suppl. Fig. 1D-F**). It is well known that 15mers more efficiently activate CD4+ T cells compared to CD8+ T cells. We therefore focused subsequent analyses on CD4+ T cells.

New Leiden clustering of CD4+ T cells yielded naïve-like, TCM-like, TEM-like, and cytotoxic T cells, in addition to regulatory T cells and cycling cells (**Fig. 1B-C; Table S5**). The reactive cluster now mainly corresponded to cluster 12, showing elevated *IFNG* expression upon *ex-vivo* restimulation (**Fig. 1D**). Markers that are independent from distinct Th lineages like *CD40LG* (data not shown) and a proliferation score (**Fig. 1E**) confirmed this assignment of the reactive cluster. We next performed a longitudinal assessment in the same donors (donor A5 shown in **Fig. 1F-I**; donor A4 shown in **Suppl. Fig. 2A-D**). This revealed that *IFNG* expression in reactive clones – defined as cells located in the reactive cluster itself or belonging to clones with sister cells in the reactive cluster – was observed only at acute time points, particularly after secondary and tertiary immunization (**Fig. 1F; Suppl. Fig. 2A**). In contrast, cells remaining in the reactive cluster at the memory time point were negative for *IFNG* (**Fig. 1F; Suppl. Fig. 2A**). The dynamics of the magnitude of reactive cells thereby resembled IFNγ ELISpot analysis for the same donor (**Fig. 1G-I; Suppl. Fig. 2B-D**).

### Validation of reactive clonotypes

We sought to validate that clonotypes recruited into the reactive cluster indeed represented SARS-CoV-2-reactive T cells. Intuitively, one would expect such clones to be recruited into the reactive cluster only in the stimulated condition, while being among unreactive cells in the unstimulated condition. Investigating the phenotype and TCR sequence of cells in the reactive cluster in the unstimulated and stimulated conditions, we noted that several clonotypes in donor A5, but not in donor A4, corresponded to mucosal-associated invariant T (MAIT) cells (**Fig. 2A; Suppl. Fig. 2E-G**). We quantified, for each clonotype, the number of cells present in the reactive cluster under stimulated or unstimulated conditions. We observed two particularly expanded clonotypes: the MAIT clone 14370 and the conventional clone 17690. Both clones were abundant in the reactive cluster, but irrespective of the stimulation condition (**Fig. 2B**). By contrast, other clonotypes, such as clone 18600, consisted exclusively of unreactive cells in the unstimulated condition but recruited most of their cells into the reactive cluster upon stimulation, thereby exhibiting the expected pattern of antigen-reactive T cells. We also identified many smaller clonotypes represented by single reactive cells under stimulation conditions (**Fig. 2B**).

**Fig. 2:**
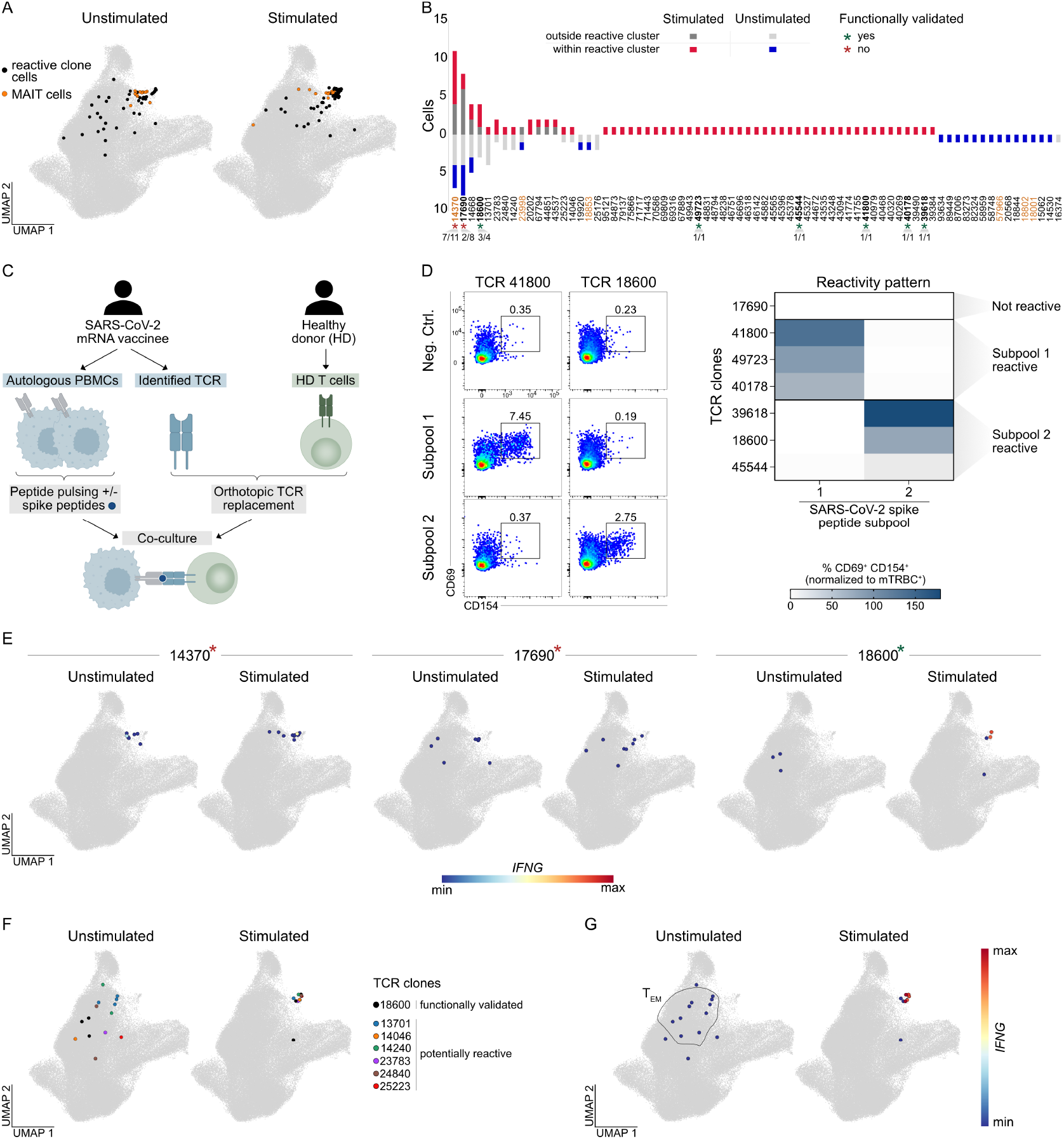
Functional validation of SARS-CoV-2 reactive T cell clones. **A** UMAP showing all cells classified as reactive (cells belonging to clones where at least one cell is in the reactive cluster) from donor A5 across all pooled time in the unstimulated (left; n=57 cells) and stimulated (right; n=83 cells) condition. Cells classified as MAIT cells (annotation described in methods section) are depicted in orange. Non-reactive cells and cells from the other donor are shown in grey. **B** Reactive clones of donor A5 are shown with the respective number of cells located within or outside the reactive cluster in the stimulated and unstimulated conditions. Clones identified as MAIT clones are highlighted in orange. Selected clones that were functionally validated and are shown in bold. For these clones, the fraction of cells recruited to the reactive cluster after re-stimulation is indicated at the bottom. Clones for which SARS-CoV-2 spike reactivity could be functionally validated are highlighted with a green asterisk, while non-reactive clones are marked by a red asterisk. **C** Experimental setup for functional validation. T cells from healthy donors (HD T cells) were equipped with TCRs identified from donor A5 by CRISPR/Cas9-mediated orthotopic TCR replacement (OTR). Transgenic T cells were co-incubated with antigen-loaded PBMCs (serving as APCs) from donor A5. PBMCs were loaded with 1 µg/mL of 15-mer peptides covering the complete wildtype spike protein (provided in two separate subpools, S1 and S2) and reactivity was assessed by flow cytometry for activation marker expression. **D** Primary data for two exemplary TCRs (left) after co-culture with SARS-CoV-2 spike peptide-loaded APCs (separately loaded with S1 and S2 peptide subpools). Data are pre-gated on living CD19^-^ CD4^+^ hTCR^-^ lymphocytes. Quantification of CD69^+^ CD154^+^ double-positive CD4^+^ T cells per clone (normalized to mTRBC^+^ cells) after SARS-CoV-2 spike-specific stimulation with S1 or S2 subpools (right). Classification of clones as non-reactive, S1-reactive, or S2-reactive is indicated on the right. Only non-MAIT clones are shown. Functional characterization of MAIT clone 14370 is provided in Suppl. Fig. 3. **E** *IFNG* expression in unstimulated or stimulated T cells for three representative clonotypes classified as reactive in donor A5. For each clonotype, cells belonging to that clonotype are shown in an individual paired panels (unstimulated condition on the left, stimulated condition on the right), while cells not belonging to that clonotype are shown in grey. Each clone is annotated with its functionally validated SARS-CoV-2 spike reactivity status (green asterisk = reactive; red asterisk = non-reactive). For the remaining functionally tested clones, see Suppl. Fig. 5A. **F-G** Based on results from Fig. 2A-E, clones with cells located outside the reactive cluster in the unstimulated and within the reactive cluster in the stimulated condition were classified as potentially reactive clones. For donor A5, cells of these clones (n=6) together with the functionally validated clone 18600 are depicted in the unstimulated (left) and stimulated condition (right) in the UMAP (F). Colors indicate cells belonging to the same clone. *IFNG* expression in unstimulated or stimulated T cells for these clones (G). Cells not belonging to these clonotypes are shown in grey. For visualization, log-transformed expression values >3 were clipped.

To functionally validate the antigen reactivity of these clonotypes, we performed CRISPR/Cas9-mediated orthotopic TCR replacement in unrelated primary human T cells from healthy donors. TCR-transgenic effector cells were co-cultured with spike antigen peptide-pulsed autologous PBMCs from SARS-CoV-2 mRNA-vaccinated donors to maintain HLA congruence (**Fig. 2C**). This approach differentiated spike-reactive TCRs from non-reactive clonotypes. While clones like 18600 or 41800 responded to sub-pools 1 or 2 of spike antigen, respectively, the expanded MAIT clone 14370 and clone 17690 both lacked reactivity (**Fig. 2D; Suppl. Fig. 3-4**). We confirmed the MAIT origin of TCR 14370 by a positive MR1 tetramer staining with the MAIT ligand 5-OP-RU next to prototypical TRAV1-2 staining (**Suppl. Fig. 3**). However, the MAIT TCR 14370 neither reacted to spike antigen nor to the HLA class I-restricted SARS-CoV-2 epitope HLA-B*15:01/NQKLIANQF, which we tested because recognition of this epitope is reported for the identical CDR3α and CDR3β chains in VDJdb^27^. Overall, TCR re-expression in primary human T cells validated spike reactivity of those non-MAIT clones that, in reverse phenotyping, showed specific recruitment to the reactive cluster upon stimulation.

Visualization of exemplary spike antigen-reactive or unreactive clonotypes in the UMAP space further demonstrated the specificity of this validation (**Fig. 2E; Suppl. Fig. 5**). Clones with unreactive TCRs – notably including MAIT clones like 14730 – showed a dispersed distribution across reactive and non-reactive clusters in both unstimulated and stimulated conditions, without specific upregulation of *IFNG* after stimulation. In contrast, validated clones like 18600 displayed both selective recruitment into the reactive cluster and robust *IFNG* induction, but only in the stimulated condition. From here on, we excluded MAIT clones from our further analyses and concentrated on clones that are likely truly reactive given a reactivity pattern resembling the one of clone 18600, thereby also excluding clone 17690 (**Fig. 2F**). Those clones were exclusively recruited into the reactive cluster upon *ex-vivo* stimulation, while without such stimulation they were mostly found in the TEM clusters 7 and 11 (**Fig. 2G**). Of note, the TEM clusters were characterized by low *TBX21* and *IFNG* expression (**Fig. 1C**), highlighting the phenotypic changes antigen-reactive clones underwent upon antigenic stimulation *in vitro*.

### Complementary validation by pHLA class II dextramers

Having established the reverse phenotyping approach for post-vaccination samples and validated antigen-reactive clonotypes, we next sought to complement our analysis by an orthogonal strategy. To this end, we used pHLA class II dextramers for the S_167_ epitope (next to various pHLA class I dextramers for spike epitopes^26^) and performed scRNAseq and scTCRseq, including a CITEseq antibody cocktail for 130 surface proteins after flow cytometric sorting of CD4+ and CD8+ T cells (**Suppl. Fig. 6A**). Assignment of cells and clones as “dextramer-positive” depended on dextramer unique molecular identifier (UMI) counts as well as cellular and clonal purity thresholds for a given dextramer (see methods). S_167_-binding clones could be unequivocally distinguished from clones binding to pHLA class I spike epitopes and were found in six out of seven tested donors, including a unique case of a hypervaccinated individual from Magdeburg (HIM) whose immune response to 217 SARS-CoV-2 vaccinations we previously characterized^28^ (**Suppl. Fig. 6B-G**).

Clustering analysis based on transcriptomes identified regulatory T cells in cluster 6, activated T cells in cluster 5, naïve-like cells in clusters 4, 0, and 2, EM-like cells in cluster 1, and cytotoxic CD4+ cells in cluster 3 (**Fig. 3A-B; Table S6**). The cytotoxic cluster mostly contained expanded clonotypes and expressed *CCL5*, whereas the EM-like cluster 1 partly maintained stemness markers such as *SELL* and *CCR7* (**Fig. 3C-D**).

**Fig. 3:**
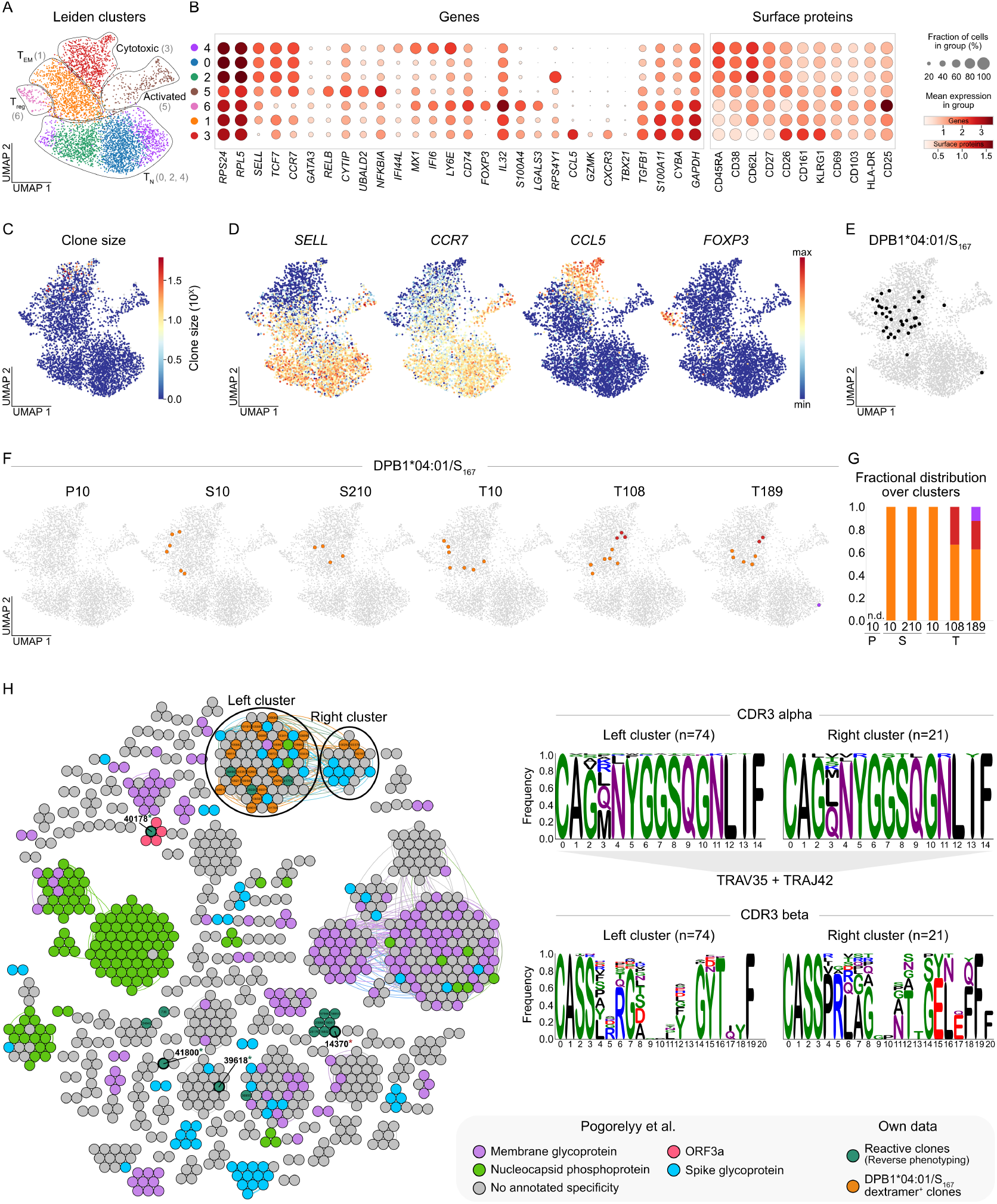
Identification of SARS-CoV-2-specific T cell responses by pHLA class II dextramers. **A** UMAP with Leiden clusters (cluster names in UMAP, cluster numbers on the right) of CD4^+^ T cells enriched for pHLA dextramer-binding (Suppl. Fig. 6A) (n=4,882 cells). **B** Dot plots showing log-normalized expression of representative genes and centered log-ratio (CLR) transformed expression of surface proteins per cluster. Numbers on the left indicate cluster numbers. **C** Visualization of clone sizes for all cells in the UMAP. **D** Log-normalized expression of selected genes and scores. **E** Visualization of SARS-CoV-2 spike epitope-specific T cells (DPB1*04:01/S167: n=35) across all HLA-matched CoVa-Adapt donors excluding A7 due to breakthrough infection (n=5) and screened time points. **F-G** UMAP visualization (F) and quantified fractional distribution (G) of DPB1*04:01/S_167_-specific T cells of HLA-matched CoVa-Adapt donors (n=6) at individual time points after primary (P), secondary (S) and tertiary (T) vaccination. Colors represent cluster location of epitope-specific cells at the respective time points. Cells without the indicated epitope-specificity are shown in grey. Time points with no detected epitope-specific cells are indicated as n.d.. **H** Similarity network of TCRs identified as reactive in the reverse phenotyping dataset (dark green) and of DPB1*04:01/S_167_ dextramer^+^ TCR clones (orange) together with previously published SARS-CoV-2-specific TCRs^32^ (left). Each vertex in the similarity network represents a unique paired αβTCR clonotype, and edges connect vertices with ≤120 TCRdist units. Only clusters containing at least two clonotypes are shown. Colors indicate SARS-CoV-2 epitope-specificity and dataset origin. Clonotypes without assigned specificity in the published dataset are shown in grey. Clone IDs from our own datasets are provided for each vertex, with functionally tested clones highlighted in bold. Clones for which SARS-CoV-2 spike reactivity could be functionally validated are highlighted with a green asterisk, while non-reactive clones are marked by a red asterisk. Two clusters containing our reactive and DPB1*04:01/S_167_-specific clones, together with spike-annotated published clones, are highlighted (left and right clusters). Sequence motifs of CDR3α and CDR3β regions for the highlighted clusters are shown on the right. Amino acid positions are indicated at the bottom of each plot. For the CDR3α region, the dominantly used V- and J-gene segments are indicated below the motifs. For detailed V- and J-gene segment usage, see Suppl. Fig. 6H.

Dextramer-positive CD4+ T cells binding the S_167_ epitope were markedly concentrated in the EM-like cluster 1, which remained stable upon longitudinal assessment (**Fig. 3E-G**). Top differentially expressed genes (DEG) in this cluster 1 (**Table S6**) included *FTH1* (ferritin heavy chain 1), previously described in SARS-CoV-2 specific CD4+ T cells with an intermediate differentiation degree^18^; *ZFP36* and *ZFP36L2*, who code for zing finger protein 36 family members that restrain full-blown T cell activation being RNA binding proteins^29^; and glycolysis/Th1 signature genes like *GAPDH, AHNAK, ANXA1* and *CYBA*^30^.

To integrate our reverse phenotyping approach with the dextramer data, we clustered antigen-reactive and dextramer-binding TCRs identified through both strategies based on sequence similarity (TCRdist^31^) together with previously published SARS-CoV-2-specific TCRs^32^, including TCRs reactive to the spike surface glycoprotein and the nucleocapsid phosphoprotein (**Fig. 3H**). This revealed two large related clusters in which TCRs co-clustered that had been identified as spike-specific by our reverse phenotyping, by our dextramer analysis, and by others before^32^. The two presumably S_167_ reactive clusters exhibited similar, yet differing sequence motifs in their complementary determining regions of the TCRβ chain (CDR3β) and a striking convergence in CDR3α (**Fig. 3H**) with a TRAV35/TRAJ42 bias (**Suppl. Fig. 6H**), which has been previously reported for TCR sequences recognizing this particular epitope^8,25,33^. This underlined the consistency of our approach and allowed us to stratify TCRs identified through reverse phenotyping as likely specific for the S_167_ epitope versus TCRs more likely directed to other regions of the spike protein.

Our integrated TCRdist analysis, powered by our validation experiments with TCR-transgenic T cells, also yielded other important observations. For example, it pointed to a presumably spike-reactive (false-positive) cluster entailing the non-validated clone 14370, highlighted alternative clusters recognizing the S_167_ epitope, including the validated clone 39618, and revealed a cluster designated as ORF3a-reactive entailing the validated spike-reactive clone 40178. Overall, this underlines the usefulness of integrating analyses with narrow epitope-specific and broad antigen-reactive resolution, and stresses the importance of reactivity validation through transgenic TCR re-expression.

## Discussion

Taken together, our data revealed transcriptional signatures in SARS-CoV-2 spike-reactive CD4+ T cells following mRNA vaccination in the absence of *ex vivo* restimulation. Our findings confirmed an expected induction of activation markers upon *in-vitro* stimulation but placed this phenotypic plasticity into the context of the unperturbed *ex vivo* phenotypes.

The “Hawthorne effect” from behavioral psychology^34^: describes how the act of measurement can itself alter the outcome. A similar concept exists in physics, where observation can influence the state of a system, as illustrated by Young’s double-slit experiment^35^ and Schrödinger’s cat thought experiment^36^. Analogously, *in-vitro* restimulation of T cells acts as an observer effect: the process of restimulation alters the cells’ phenotype, and the specific conditions of restimulation shape the resulting state. One might argue that a cell’s functional state – what it does – is more relevant than its resting phenotype – what it looks like. In human immunology, however, it is easier to describe the *ex-vivo* phenotype compared to the *in-vivo* function. This discrepancy becomes important when the same T cell can display different activation profiles depending on the context.

For example, there is a long-standing discussion on how to define Th1 cells^37^. A functional definition classifies Th1 cells as those producing IFNγ upon stimulation. A transcriptional definition highlights expression of the lineage-defining transcription factor T-bet (encoded by *TBX21*). If functional evaluation is not feasible, combinations of chemokine receptors are sometimes used as surrogates, in case of Th1, for example, CXCR3^37^. While it would be correct to classify spike-reactive CD4+ T cells as Th1 cells because they upregulate *IFNG* transcripts upon stimulation, in the resting state they rather showed a “Th-neutral” phenotype, indicating plasticity depending on the stimulation conditions.

This has important implications for single-cell sequencing studies, which usually analyze cells in a resting state without restimulation. It is critical to recognize that such analyses capture a baseline phenotype and that transcription factor expression may not reflect what the same cells would look like after stimulation. The same caveat applies to *ex vivo* analyses using pHLA tetramers or dextramers, whether by flow cytometry or single-cell sequencing. The strength of such approaches is that stimulation is not required, but the limitation is that the resulting phenotype will differ from that observed after *in vitro* stimulation.

Several limitations of our study need to be acknowledged. In the unstimulated condition, we defined reactive clonotypes as those with cells in the reactive cluster, or as unstimulated clonotypes with sister cells in the reactive cluster in the stimulated condition. This definition introduces a potential bias toward clones detectable in the reactive cluster. Conceivably, there may be small clonotypes that never enter the reactive cluster in either condition. Such clonotypes would not be categorized as reactive by our approach. However, their relevance is likely limited, since they are neither abundant nor show disproportionate recruitment upon stimulation.

Another consideration is the number of donors available for reverse phenotyping. While our study applies this method in a standardized longitudinal setting for the first time, reverse phenotyping is highly cost-intensive. Large numbers of cells must be sequenced by scRNAseq across both stimulated and unstimulated conditions, without pre-enrichment for antigen reactivity. Although we pre-enriched for non-naïve T cells, thereby potentially excluding naïve-like or stem cell memory CD4+ T cells, deep unbiased sequencing was still required to capture rare reactive clones with limited cell numbers. For this reason, we do not view reverse phenotyping as a technique for routine donor characterization. Instead, we see it as a screening method for hypothesis generation to highlight the phenotypic biases introduced by restimulation and to reveal otherwise hidden states in the unstimulated condition. Subsequent targeted approaches, such as dextramer-based and functional assays, can then be used for more efficient analysis, including assessment of molecular candidates on the protein level.

A strength of our study is the validation of TCRs after re-expression in primary human T cells. This revealed that some expanded clonotypes, including MAIT cells, were nonspecifically present in the reactive cluster under both stimulated and unstimulated conditions. Single-cell transcriptomes had already suggested that these were unlikely to represent true antigen-reactive CD4+ T cells, since they did not show selective recruitment upon stimulation. TCR re-expression confirmed this suspicion by demonstrating a lack of spike reactivity. Without reverse phenotyping, such bystander clonotypes would likely have been misclassified as reactive. For example, had we simply stimulated PBMCs and sequenced all cells expressing activation markers or IFNγ, we would have included many of these expanded bystanders. Large clones may be particularly prone to this effect, as their size makes it more likely to sample cells in diverse activation or differentiation states.

In summary, our study demonstrates that reverse phenotyping combined with clonotypic tracking and functional TCR validation provides a powerful framework to dissect the biology of antigen-reactive T cells. By comparing stimulated and unstimulated states, we highlight the extent to which *in vitro* restimulation reshapes T cell phenotypes and how resting transcriptional states can nevertheless preserve a memory of past antigen encounter. These insights have direct implications for the interpretation of single-cell studies and for the classification of T cell subsets, reminding us that definitions based solely on either function or phenotype may be incomplete. More broadly, our work emphasizes the value of integrating orthogonal technologies – from unbiased transcriptomics to epitope-specific dextramers and functional TCR assays – to obtain a more accurate and nuanced understanding of human T cell immunity after vaccination^38^.

## Methods

### Study design

This was an exploratory study; therefore, a formal power analysis was not feasible. The sample size was predefined based on our experience and that of others in similar research contexts. The primary rationale of the study was to investigate the immunobiological mechanisms of adaptive immune responses following immunization, specifically with SARS-CoV-2 mRNA vaccines. No randomization or blinding was performed. The number of experimental replicates is indicated at the relevant sections of the manuscript. Blood samples from healthy donors who received a vaccination at the Vaccination and Travel Clinic of the University Hospital Erlangen were used as study material and served as the inclusion criterion. The following individuals were excluded from participation in this study: individuals unable to provide informed consent, minors, pregnant or breastfeeding individuals, and persons in a dependent relationship with other study participants or the study investigators.

### Study cohort CoVa-Adapt

Ethics approval was granted by the local Ethics Committee of the Medical Faculty of the University Hospital of Erlangen, Friedrich-Alexander University Erlangen-Nürnberg, Germany (350_20B). Samples were collected after informed consent of the donors. Analyses from this cohort have been previously published^26,28,39^ and the cohort (DRKS-ID: DRKS00034356) is described in detail in Table S1. HLA-typing was previously published^26^. In brief, donors were 23-58 years old (median: 42; interquartile range (IQR) 29-51), 55% female, of European Caucasian ethnicity, overall healthy (no chronic medication), of normal weight, and received a three-dose mRNA vaccination regimen with Comirnaty. Time intervals between immunizations were 22-28 days (median 23, IQR 23-25) between 1^st^ and 2^nd^ dose, and 196-244 days (median 227, IQR 216.5-229) between 2^nd^ and 3^rd^ dose. Blood was sampled before the 1^st^ dose, 8-13 days post 1^st^ dose (median 10, IQR 9-10), 9-11 days post 2^nd^ dose (median 10, IQR 10-11), 195-211 days post 2^nd^ dose (median 210, IQR 209-210), 9-12 days post 3^rd^ dose (median 10, IQR 10-10) and 157-202 days post 3^rd^ dose (median 189, IQR 189-190). Additional blood samples were collected for seven donors 101-126 days (median 108, IQR 108-113.5) after 3^rd^ dose. Some donors experienced SARS-CoV-2 breakthrough infections after day 10 post 3^rd^ dose which is indicated in the figures or figure legends if applicable. These breakthrough infections were self-reported and confirmed by PCR tests. 44% of breakthrough infections could be additionally validated by SARS-CoV-2 nucleocapsid^+^ serology, while no infection-free donor had detectable SARS-CoV-2 nucleocapsid-specific antibodies.

### Peripheral blood mononuclear cell (PBMC) isolation

PBMCs were isolated from citrated peripheral blood by density gradient centrifugation using a BioColl density medium with a density of 1.077 g/mL (BioSell, BS.L 6115). Cells were resuspended in heat-inactivated FCS + 10% DMSO and stored in liquid nitrogen.

### Multiparametric flow cytometry for T cells

The following antibodies were used for T cell analysis: from BD Biosciences: anti-CD4-BUV395 (564724, 1:200), anti-CD19-PE/CF594 (562294, 1:200), anti-CD69-PerCP-Cy5.5 (560738,1:100); from BioLegend: anti-CD4-BV510 (300545, 1:50), anti-CD8-APC (301049, 1:400), anti-human a/b TCR (hTCR)-FITC (306706, 1:200), anti-mouse TCR β chain (mTRBC)-APC/Fire™750 (109246, 1:100), anti-CD45RA-PerCP-Cy5.5 (304121,1:400), anti-CD62L-PacificBlue (304825,1:400); from eBioscience: anti-CD4-PE (12-0049-42, 1:400), anti-CD8-eF450 (48-0086-42, 1:200), anti-CD56-FITC (11-0566-42, 1:200), anti-CD45-PerCP-Cy5.5 (45-0459-42, 1:100), anti-CD45-PE/Cy7 (25-9459-42, 1:400); from Dako: anti-CD45-PacificBlue (PB986, 1:50); from Miltenyi: anti-CD154-APC (130-113-603,1:100). For the viability staining, ethidium-monoazide-bromide (EMA; ThermoFisher, E1374), propidium iodide (PI; ThermoFisher, P1304MP), or ZombieNIR Fixable Viability Kit (BioLegend, 423106, 1:1000) was used.

Cells were first washed twice with cold FACS buffer (PBS + 0.5% BSA). For experiments involving fixation and permeabilization, cells were resuspended in 100 µL of FACS buffer containing EMA (1:1000), incubated under bright light for 15 min, and washed once with cold FACS buffer before being incubated with 25 µL of surface antibody mix. In experiments without fixation/permeabilization, cells were directly resuspended in 25 µL of surface antibody mix and incubated on ice for 20 min. Surface staining was followed by two washes with cold FACS buffer if EMA staining had been performed, or one wash if not. In the latter case, cells were subsequently resuspended in 50 µL of FACS buffer containing PI (1:100), incubated for 3 minutes, and washed twice with cold FACS buffer. Where applicable, fixation and permeabilization were performed using the BD Cytofix/Cytoperm Kit (BD Biosciences, 554714) according to the manufacturer’s instructions, followed by one wash with cold Perm/Wash buffer (1×) and one final wash with cold FACS buffer. Samples were acquired on a LSRFortessa Cell Analyzer (BD Biosciences) and analyzed with FlowJo 10.7.2 (Tree Star Inc.).

#### IFNg Enzyme-linked immunospot (ELISpot)

Cryopreserved PBMCs were thawed and rested overnight at 1×10^6^ cells/mL in complete RPMI medium (cRPMI: RPMI 1640 Medium + 10% heat-inactivated FCS, 0.05 mM β-mercaptoethanol, 0.05 mg/mL gentamicin, 1.1915 g/L HEPES, 0.2 g/L L-glutamine, 100 U/mL Penicillin-Streptomycin). ELISpot plates (Millipore, MSIPS4510) were coated with anti-human IFNg monoclonal antibody (clone 1-DIK, Mabtech, 3420-3-1000) at 0.5 µg/well overnight at 4°C. Plates were washed with sterile PBS and subsequently blocked with cRPMI medium for 1-2 h at 37°C. PBMCs were seeded at a density of 400,000 cells/well and stimulated with 11aa overlapping 15-mer PepMix™ SARS-CoV-2 spike glycoprotein peptide pool (JPT, PM-WCPV-S-2) or SARS-CoV-2 nucleoprotein peptide pool (JPT, PM-WCPV-NCAP-2) at a concentration of 1 µg/mL for 20 h at 37°C. For the unstimulated condition, PBMCs were cultured in cRPMI medium and respective dilution of solvent DMSO. As a positive control, PBMCs were stimulated with 25 ng/mL phorbol myristate acetate (PMA) (Sigma-Aldrich, P1585-1mg) and 1 µg/mL ionomycin (Sigma-Aldrich, I9657-1MG). The following steps were performed at room temperature. Plates were washed with PBS containing 0.05% Tween-20 (Sigma-Aldrich, P9416-50ml) and incubated with biotinylated anti-human IFNγ monoclonal antibody (clone 7-B6-1, Mabtech, 3420-6-250) at 0.2 µg/well for 2 h. Plates were washed a second time with PBS containing 0.05% Tween-20 and subsequently incubated with an avidin-biotinylated peroxidase complex (VECTASTAIN^®^ Elite ABC-HRP Kit, Vector Laboratories, VEC-PK-6100) for 1-2 h. Afterwards, plates were washed first with PBS containing 0.05% Tween-20 following one washing step with PBS. Plates were developed by the addition of AEC substrate solution (Sigma-Aldrich, 152224-10mL) for 15 min, washed with water and dried for 24 h in the dark. Analysis was performed on an ImmunoSpot^®^ Analyzer (Cellular Technologies Limited).

#### Reverse phenotyping

Single-cell RNA sequencing (scRNAseq) was performed for PBMCs from two CoVa-Adapt donors at a total of four time-points after primary (P), secondary (S), and tertiary (T) vaccination. Three independent experiments (Experiments 1–3) were conducted and pooled into a single dataset (see Table S2). PBMCs were thawed from cryopreserved stocks and rested overnight at 1×10^6^ cells/mL in cRPMI medium.

For each donor, 2×10^6^ PBMCs were stimulated with stimulated with 11aa overlapping 15-mer PepMix™ SARS-CoV-2 spike glycoprotein peptide pool (JPT, PM-WCPV-S-2) at a concentration of 1 µg/mL for 4 h at 37°C. For the unstimulated condition, PBMCs were cultured in cRPMI medium and respective dilution of solvent DMSO. After stimulation, cells were washed with cold FACS buffer and stained with surface antibodies (anti-CD19-PE/CF594, anti-CD56-FITC, anti-CD8-APC, anti-CD4-PE, anti-CD62L-PacificBlue, anti-CD45RA-PerCP-Cy5.5). In Experiment 3, individual hashtag antibodies (2.5 µL per 5×10^6^ PBMCs of TotalSeq-C anti-human hashtag antibodies 1–8, BioLegend, 394661, 394663, 394665, 394667, 394669, 394671, 394673, 394675) were included for sample multiplexing. All samples were incubated for 30 minutes on ice, followed by two washes with cold FACS buffer. Live/dead discrimination was performed using PI (1:200) immediately before sorting. Single, live, CD19- and CD56-negative, CD4- or CD8-positive, non-naïve (defined as CD45RA-positive and CD62L-negative, CD45RA-negative and CD62L-negative, or CD45RA-negative and CD62L-positive) lymphocytes were sorted in previously FCS-coated 96-well U-bottom plates filled with FACS buffer. In Experiment 3, eight different samples were pooled prior to sequencing and later distinguished using the unique TotalSeq-C hashtags during scRNAseq analysis. Cells were sorted on a FACS Aria II cell sorter (BD).

Immediately after sorting, cells were loaded to a Chromium Next GEM Chip K (10X Genomics) and Chromium Next GEM Single-Cell 5′ kits v.2 were used to generate GEX, VDJ and CITEseq libraries according to the manufacturer’s instructions (10X Genomics, 1000263, 1000256, 1000252, 1000286, 1000250, 1000215, 1000190). Libraries were sent to Novogene (Cambridge, UK) and sequenced on an Illumina NovaSeq platform with PE150 strategy.

#### Dextramer staining

Single-cell RNA sequencing was performed for PBMCs from six CoVa-Adapt donors at a total of six time-points after primary (P), secondary (S), and tertiary (T) vaccination as well as for PBMCs from HIM 189 days after 215th vaccination (see Table S3). PBMCs were thawed from cryopreserved stocks and rested overnight at 1×10^6^ cells/mL in cRPMI medium.

For each donor, individual dextramer cocktails were prepared directly before staining (see Table S3). For each cocktail, 1 µL per 5×10^6^ cells of HLA-matched dCODE Dextramers^®^ (immudex) and 100 µM d-biotin (1/10 of total dextramer volume) were pre-mixed in FACS-buffer to block free binding sites. 5×10^6^ PBMCs per donor and time-point were recovered and first stained with 50 µL of individual dextramer cocktails for 30 min on ice. Afterwards, a cocktail of surface antibodies and viability staining (anti-CD19-PE/CF594, anti-CD56-FITC, anti-CD8-APC, anti-CD4-BV510, ZombieNIR), individual anti-CD45 antibodies (anti-CD45-PacificBlue, anti-CD45-PerCP/Cy5.5, anti-CD45-PE/Cy7), individual hashtag antibodies (2.5 µL per 5×10^6^ PBMCs of TotalSeq-C anti-human hashtag antibodies 1–8, BioLegend, 394661, 394663, 394665, 394667, 394669, 394671, 394673, 394675), and TotalSeq-C antibodies (0.078 µg per 5×10^6^ PBMCs anti-human CD45RA (BioLegend, 304163), 0.277 µg per 5×10^6^ PBMCs anti-human CCR7 (BioLegend, 353251), 0.25 µg 5×10^6^ PBMCs anti-human CXCR3 (BioLegend, 353251)) were added. For three samples, instead of individual TotalSeq-C antibodies, the TotalSeq-C Human Universal Cocktail, V1.0 (BioLegend, 399905) was added. On the day of the experiment, three vials of this universal cocktail were rehydrated in 18 µL of FACS buffer each, incubated for 5 min at room temperature, vortexed, and centrifuged at maximum speed for 30 sec at room temperature. One vial was used to stain 5×10^6^ PBMCs. Samples were incubated for an additional 30 min on ice. Cells were washed four times with cold FACS buffer and up to 8 samples were pooled prior to the sort. Pooled samples could be distinguished by individual CD45 color-barcoding at the sorter and by individual TotalSeq-C anti-human hashtag antibodies in the scRNAseq dataset. Single, live, CD19- and CD56-negative, CD4- or CD8-positive, dextramer-positive lymphocytes were sorted in previously FCS-coated 1.5 mL tubes filled with FACS buffer. Additionally, single, live, CD19- and CD56-negative, total CD4- or CD8-positive lymphocytes were sorted from three donors, as a framework for CD4- and CD8-T-cell phenotypic clusters. Cells were sorted on a FACS Aria II cell sorter (BD).

Immediately after sorting, cells were loaded to a Chromium Next GEM Chip K (10X Genomics) and Chromium Next GEM Single-Cell 5′ kits v.2 were used to generate GEX, VDJ and CITEseq libraries according to the manufacturer’s instructions (10X Genomics, 1000263, 1000256, 1000252, 1000286, 1000250, 1000215, 1000190). Libraries were sent to Novogene (Cambridge, UK) and sequenced on an Illumina NovaSeq platform with PE150 strategy.

### Computational single-cell RNA sequencing data

#### analysis

The Cellranger (version 6.0.2, 10x Genomics) ‘multi’-command was used to process the individual sequencing runs of the reverse phenotyping and the dextramer dataset using the references GRCh38 (version 2020A, 10x Genomics) and vdj-GRCh38 (version 5.0.0, 10x Genomics). For sequencing runs that include dextramers, CITE-markers, or hashtag antibodies, additional customized feature barcode references were used. The remaining analysis was performed using Scanpy (v.1.8.2^40^) and Scirpy (v.0.10.1^41^) following recent best practices^42^.

#### Quality Control

GEX, antibody capture, and TCR data were fused on the run-level. Low-quality cells, such as doublets and dying cells, were filtered through thresholds on UMI counts, the number of detected genes, and the fraction of mitochondrial genes. Genes infrequently expressed in less than 10 cells were excluded. The gene expression count matrices were log1p-transformed after normalization to 10,000 counts per cell. Cells without TCR and outlier clusters were excluded from further analysis.

#### Annotation

Annotation was performed on a dataset level after fusing all runs from the reverse phenotyping and the dextramer dataset independently. Samples including hashtag antibodies were annotated for donor and time points via HashSolo^43^, and any annotated doublets or negative cells were removed. Cells with identical CDR3α and CDR3β amino acid sequences on primary or secondary chains were annotated as individual clonotypes, and clonal expansion was calculated per donor. For comparability, the IDs of clonotypes occurring in both datasets were unified. Scores of represented marker genes were calculated on a cell-level as described in their corresponding publication^2,44^. MAIT cells were annotated by the CDR3α sequence CAVMDSSYKLIF or by the combination of TRAJ33, TRAV1-2 together with TRBV20-1 or TRBV6 TCR genes. CD4^+^, CD8^+^, double positive, and double negative T cells were defined by the combination of corresponding marker genes CD4, CD8A, or CD8B at a threshold of 0.5. When protein surface markers were available, this was complemented by Hu.CD8 at 0.75 and Hu.CD4_RPA.T4 at 0.95. If not stated otherwise, only CD4^+^ cells were considered for the analysis. In the reverse phenotyping dataset, γδ cells were defined at a γδ-marker score greater than 0 and not expressing a TCR. All cells with TCR and a γδ-score less than 0 were considered αβ cells. Leiden clustering^45^ (resolutions: *r*_*RP_full*_*=0*.*75, r*_*RP_CD4*_*=1*.*25, r*_*RP_unstim*_*=0*.*75, r*_*Dex_CD4*_*=0*.*8*) and UMAP visualisation^46^ (*n*_*neighbors*_*=15*) were performed on the 5,000 most variable genes without TCR-forming genes on various subsets of the datasets.

#### Reactivity and specificity

In the reverse phenotyping dataset, the reactive cluster was identified by elevated INFG and Proliferation score levels, as well as a high fraction of stimulated cells. All cells of a reactive clonotype with at least one member in the reactive cluster were marked as reactive. In the dextramer dataset, specificity towards TFEYVSQPFLMDLE was assigned for cells with a UMI count of 6 or higher. Additionally, the counts for TFEYVSQPFLMDLE were required to amount to at least 40% of total UMIs, and 50% of the clonotype members must have expressed specificity.

#### Analysis

Differential expression analysis of genes and surface proteins was performed via ‘scanpy.tl.rank_genes_groups’ through a t-test with Benjamini-Hochberg correction. For analysis on clonotype similarity, the reverse phenotyping dataset and the dextramer dataset were combined with annotated TCR data from Pogorelyy et al.^32^ TCR clonotypes were clustered using TCRdist3 as a distance measure^31^. Connected components of clonotypes within 120 TCRdist units were formed, and visualized through the ‘Circle Pack Layout’ in Gephi (v.0.9.7^47^) with component ID and modularity as hierarchy. CDR3 motifs were calculated on the respective amino acid sequence aligned by MUSCLE^48^ and visualized through Logomaker^49^.

### Data and Code Availability

The single-cell sequencing data are publicly available at NCBI GEO under the accession numbers GSE310441 and GSE310442 for the reverse phenotyping dataset as well as GSE249998 for the dextramer dataset. The annotated data used for analysis is available on Zenodo under the accession 10.5281/zenodo.17091562. Single-cell analysis code: https://github.com/SchubertLab/CovidVac_CD4.

### Transgenic TCR re-expression in primary human T cells

A coherent description for the workflow of targeted TCR re-expression in primary human T cells using CRISPR-Cas9-mediated orthotopic TCR replacement has previously been published^26,50,51^. A brief description including all relevant alterations to the published protocol is summarized in the following chapters ‘Transgenic TCR DNA template design’, ‘Double-stranded DNA production’, ‘T cell activation for genetic editing’, ‘Ribonucleoprotein production’, and ‘Orthotopic TCR replacement (OTR)’.

### Transgenic TCR DNA template design

The DNA template was designed *in silico* and synthesized by Twist Bioscience. The construct had the following structure: The left homology arm (LHA; 396 bp) was followed by a self-cleaving peptide P2A and the TCR β-chain which consisted of the human variable part (VDJβ) and the murine TCR β constant region with an additional cysteine bridge (mTRBC-Cys)^52^. The subsequent self-cleaving peptide T2A separated the β-chain from the following a-chain which was designed according to the same principle with the human variable part (VJa) being followed by the murine TCR α constant region with an additional cysteine bridge (mTRAC-Cys)^52^. After the stop codon (TGA) and the bovine growth hormone polyA signal (bGHpA), the 330 bp right homology arm (RHA) concluded the HDR template. For sequences of these segments see **Table S7-8**.

### Double-stranded DNA production

The DNA construct was ordered as a sequence-verified plasmid gene via a commercial provider (Twist Bioscience). The lyophilized plasmid was reconstituted with sterile water to 60 ng/µL and amplified by PCR to generate a linearized double-stranded HDR template. A truncated Cas9 Target Sequences (CTS) was incorporated at the 5’-end of the HDR template by the genomic primer targeting the hTRAC LHA. Each 100 µL PCR reaction contained 1 x Herculase II Reaction Buffer, 0.4 µM hTRAC HDR genomic forward primer targeting LHA (5’-TCTCTCTCTCAGCTGGTACACGGCTGCCTTTACTCTGCCAGAG-3’), 0.4 µM hTRAC HDR genomic reverse primer targeting RHA (5’-CATCATTGACCAGAGCTCTG-3’), 0.5 mM dNTPs, 1 µL Herculase II Fusion DNA Polymerase, and 60 ng reconstituted DNA in PCR grade water. The PCR was run with the following cycling conditions: Initial denaturation at 95°C for 3 min, 34 cycles of 95°C for 30 sec, 63°C for 30 sec and 72°C for 3 min, final elongation at 72°C for 3 min, and hold at 4°C. Successful amplification was confirmed with an 1% agarose gel and amplified HDR template was purified with a MinElute PCR Purification Kit (Qiagen, 28004) according to the manufacturer’s instructions.

### T cell activation for genetic editing

PBMCs were isolated from blood provided by healthy volunteers (Transfusion Medicine, University Hospital Erlangen) and frozen at −80°C for storage. Ethics approval was granted by the local Ethics Committee of the Medical Faculty of the University Hospital of Erlangen, Friedrich-Alexander University Erlangen-Nürnberg, Germany (392_20Bc). Samples were collected after informed consent of the donors. For T cell activation, these PBMCs were thawed and overnight-rested at 2×10^6^ cells/mL in cRPMI medium supplemented with 50 U/mL Interleukin-2 (IL 2). Afterwards, PBMCs were activated for two days at 1×10^6^ cells/mL on tissue-culture flasks with 1 µg/µL of surface-bound anti-CD3 (BioLegend, 317302) and anti-CD28-antibodies (BioLegend, 302902) in medium supplemented with 300 U/mL IL 2 (Peprotech, 200-02), 5 ng/mL Interleukin-7 (Peprotech, 200-07), and 5 ng/mL Interleukin-15 (Peprotech, 200-15).

### Ribonucleoprotein production

Activated PBMCs were electroporated with ribonucleoproteins (RNPs) targeting the endogenous *hTRAC* and *hTRBC* locus as well as the purified HDR template. For final electroporation, 3.5 µL of *hTRAC* and 3 µL of *hTRBC* RNP (final concentration 20 µM) were required per electroporation sample. First, 40 µM gRNAs were produced by mixing equimolar amounts of trans-activating crRNA (tracrRNA) (Integrated DNA Technologies, 1072534) with *hTRAC*^53^ crRNA (5’-AGAGTCTCTCAGCTGGTACA-3’, Integrated DNA Technologies) or *hTRBC* crRNA^50^ (5’-GGAGAATGACGAGTGGACCC-3’, Integrated DNA Technologies) and incubating the mixtures at 95°C for 5 min with subsequent cool down to room temperature. Afterwards, 50 µg/sample poly-L-glutamic acid (PGA; Sigma-Aldrich, P4761) were added to *hTRAC* gRNA^54,55^ and 20 µM electroporation enhancer (Integrated DNA Technologies, 10007805) were added to both *hTRAC* and *hTRBC* gRNA. RNP production was concluded by adding equal volume of Cas9 Nuclease V3 (Integrated DNA Technologies, 1081059, diluted to 6 µM) to *hTRAC* and *hTRBC* gRNA (40 µM) respectively. RNPs were incubated for 15 min at RT and subsequently stored on ice for processing at the same day. For the calculations above, the volume of PGA was not considered.

### Orthotopic TCR replacement (OTR)

Prior to electroporation, DNA-sensing inhibitors RU.521 (small-molecule inhibitor of cyclic GMP-AMP synthase (cGAS); InvivoGen, inh-ru521)^54^ was added to the cells at a final concentration of 4.82 nM for 6 h. Afterwards, activation was stopped by transferring cells to a new plate in fresh cRPMI medium. For electroporation, 1×10^6^ activated cells per electroporation sample were resuspended in 20 µL P3 electroporation buffer (Lonza, V4SP-3960) and then mixed with DNA/RNP mix (0.5 µg HDR template, 3.5 µL *hTRAC* and 3 µL *hTRBC* RNPs). After transfer into the 16-well Nucleocuvette™ Strip (Lonza, V4SP-3960), cells were electroporated (pulse sequence EH100) in the Lonza 4D-Nucleofector™. After electroporation, cells were rescued in 900 µL of antibiotic-free RPMI medium supplemented with 180 U/mL IL-2. After 15 min, 100 µL of a mixture containing 0.5 µM HDAC class I/II Inhibitor Trichostatin A (AbMole, M1753) and 10 µM DNA-dependent protein kinase (DNA-PK) inhibitor M3814 (chemietek, CT-M3814) was added to each sample^56^. Cells were incubated for 12-18 h in a 24-well plate, before the medium was supplemented with an antibiotic mix containing gentamicin, penicillin and streptomycin to produce cRPMI medium. 24 h after electroporation, cells were transferred into a new 24-well plate and cultivated in a final volume of 1 mL fresh cRPMI medium supplemented with 180 U/mL IL-2. Four days after electroporation, successful editing was validated by flow cytometry (anti-CD4-PE, anti-CD8-eF450, anti-human a/b TCR (hTCR)-FITC, anti-mouse TCR β chain (mTRBC)-APC/Fire™750). Cells were cultivated for additional seven to eight days in cRPMI with addition of 50 U/mL IL-2 every two days before specificity was determined by peptide-induced activation marker expression.

### Peptide-pulsing of autologous PBMCs

To assess the reactivity of TCR-engineered T cells against SARS-CoV-2 spike epitopes, TCR-engineered T cells were co-cultured with peptide-pulsed autologous PBMCs. Autologous PBMCs were isolated and cryopreserved in liquid nitrogen as described above. One day prior to the experiment, PBMCs were thawed and rested overnight at 1×10^6^ cells/mL in cRPMI. For peptide pulsing, 1×10^6^ PBMCs per condition were incubated with 1 µg/mL of an 11aa overlapping 15-mer PepMix™ SARS-CoV-2 spike glycoprotein peptide pool or 10^-4^ M of NQKLIANQF peptide for 2 h at 37°C. After incubation, excess peptide was removed by washing. As a negative control, PBMCs were treated with an equivalent dilution of DMSO, the solvent used for the peptide pool.

### Peptide-induced activation marker expression by TCR-engineered T cells

The percentage of transgenic TCR (mTRBC)+ cells within the TCR-engineered T cell population was determined by flow cytometry one day prior to co-culture. A total of 2×10^4^ transgenic TCR (mTRBC)+ T cells were co-cultured with 2×10^4^ peptide-pulsed autologous PBMCs for 4 h at 37°C in cRPMI, in the presence of 1 µg/mL anti-CD40 blocking antibody (Miltenyi Biotec, 130-094-133). As a positive control, TCR-engineered T cells were stimulated with 25 ng/mL PMA (Sigma-Aldrich, P1585-1mg) and 1 µg/mL ionomycin (Sigma-Aldrich, I9657-1MG). Following co-culture, cells were first stained with EMA, followed by surface staining (anti-CD4-BUV395, anti-CD8-eF450, anti-CD19-PE/CF594, anti-human TCR a/b-FITC, anti-human TCR a/b-APC/Fire™750, anti-CD154-APC, anti-CD69-PerCP-Cy5.5), then fixed. In the case of MAIT TCR clone 14370, cells were first stained with EMA, followed by surface staining (anti-CD4-BUV395, anti-CD8-eF450, anti-CD19-PE/CF594, anti-human TCR a/b-FITC, anti-human TCR a/b-APC/Fire™750, anti-human TCR Va7.2-BV711), fixation and intracellular staining (anti-IFNg-FITC, anti-IL-2-APC, anti-CD154-PerCP-Cy5.5, anti-CD137-PE-Cy7). Samples were acquired on a LSRFortessa Cell Analyzer (BD Biosciences) and analyzed with FlowJo 10.7.2 (Tree Star Inc.).

### MR1 tetramer staining

MR1 tetramers loaded with the MAIT cell ligand 5-OP-RU were kindly provided by Prof. Dr. med. Jochen Mattner (University Hospital Erlangen). TCR-engineered T cells were washed twice with cold FACS buffer prior to staining. Subsequently, 25 µL of MR1 tetramers were added to each sample, and cells were incubated for 40 min at room temperature in the dark. After staining, cells were centrifuged at 480 × *g* for 5 min, and the supernatant was discarded. Cells were then stained with EMA, followed by surface staining (anti-CD4-BUV395, anti-CD8-eF450, anti-CD19-PE/CF594, anti-human TCR a/b-FITC, anti-human TCR a/b-APC/Fire™750, anti-human TCR Va7.2-BV711). Samples were acquired on a LSRFortessa Cell Analyzer (BD Biosciences) and analyzed with FlowJo 10.7.2 (Tree Star Inc.).

### Statistical analysis

Statistical analyses were performed as appropriate for each individual experiment and type of data. Specific details regarding the statistical methods used are provided in the corresponding figure legends and relevant methods section.

## Author contributions

Conceptualization: K.S., data curation: K.K., F.D., formal analysis: K.K., F.D., funding acquisition: K.S., B.Sc. investigation: K.K., F.D., C.S., B.Sp., methodology: K.K., F.D., K.S., project administration: K.S., resources: B.Sc., K.S., software: F.D., B.Sc., supervision: B.Sc., K.S., validation: K.K., F.D., visualization: K.K., F.D., K.S., writing original draft: K.K., F.D., K.S., review & editing: all authors.

## Acknowledgments

This work is supported by grants from the German Federal Ministry of Education, Technology and Space (BMFTR, projects 01KI2013 and 031L0290B), the Else Kröner-Fresenius-Stiftung (project 2020_EKEA.127) and the German Research Foundation (DFG) through the research training group RTG 2504 (project 401821119) to K.S.; and the BMFTR (project 031L0290A) to B.Sc.. F.D. is supported by the Helmholtz Association under the joint research school “Munich School for Data Science - MUDS” and acknowledges financial support from the Joachim Herz Stiftung. Funding agencies had no influence on the study design or implementation.

We thank members from the Schober laboratory for experimental support and critical discussion. We also thank Lucie Loyal (Berlin) and Petra Bacher (Kiel) for critical review of the manuscript. Furthermore, we also gratefully acknowledge the generous support by the Manfred Roth-Stiftung, Fürth, Germany, and thank the Core Unit Cell Sorting and Immunomonitoring Erlangen.

