## Supplementary Figures for "Antigen-reactive CD4+ T cells after SARS-CoV-2 vaccination show divergent phenotypic states with or without restimulation bias"

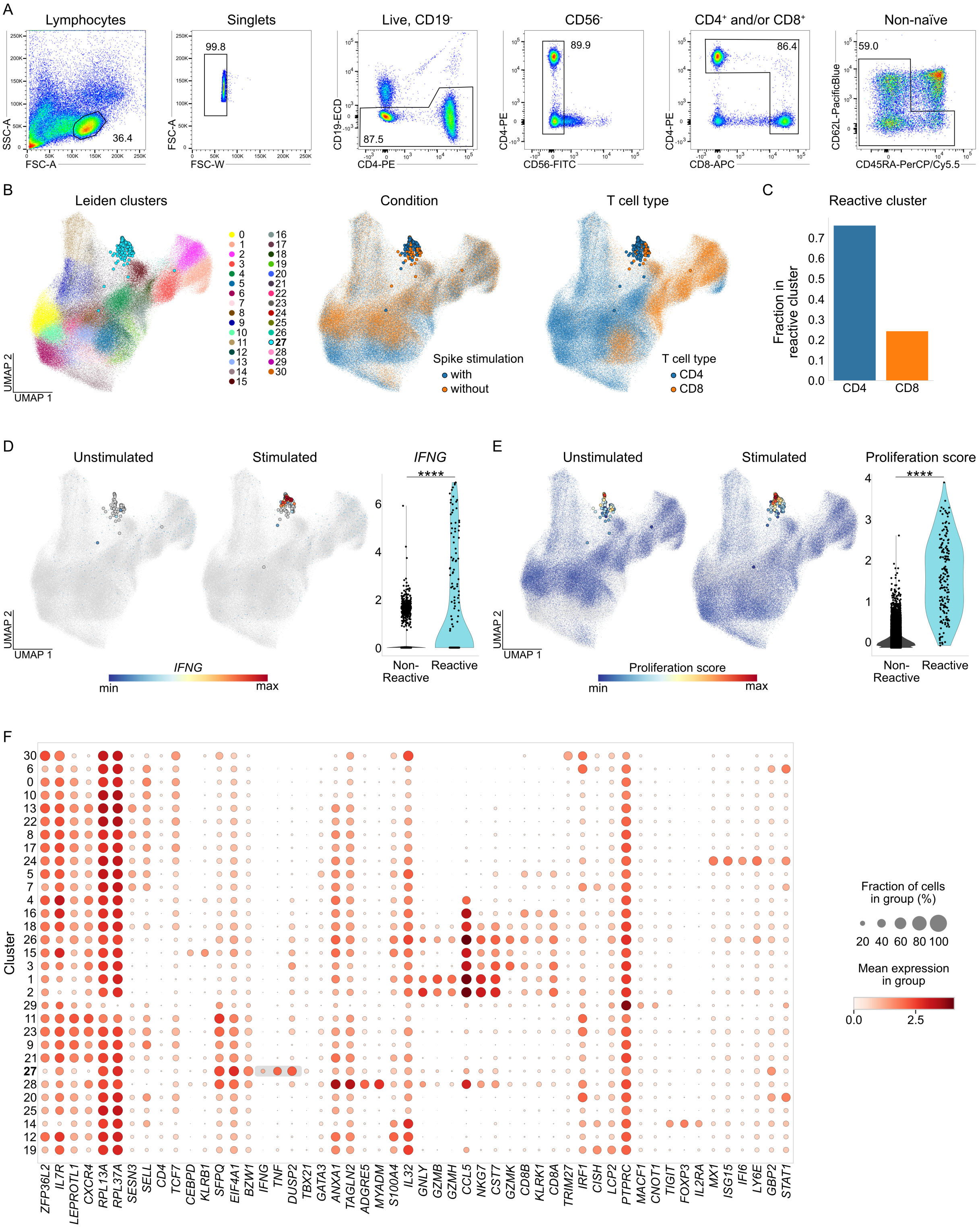


**Suppl. Fig. 1: Identification of SARS-CoV-2 spike-reactive CD4+ and CD8+ T cells via reverse phenotyping.**

*Legend see next page*

**Suppl. Fig. 1: Identification of SARS-CoV-2 spike-reactive $CD4^+$ and $CD8^+$ T cells via reverse phenotyping. A** Representative gating strategy to enrich for non-naïve $CD4^+$ and/or $CD8^+$ T cells for scRNAseq via flow cytometric cell sorting. For reverse phenotyping, PBMCs were re-stimulated with 15-mer peptides covering the complete wildtype spike protein or left untreated. Single, live, $CD19^-$, $CD56^-$, $CD4^+$ and/or $CD8^+$, non-naïve (defined as $CD45RA^-$ $CD62L^+$, $CD45RA^-$ $CD62L^-$, or $CD45RA^+$ $CD62L^-$) lymphocytes were enriched for a total of two donors across four time points after primary, secondary, and tertiary SARS-CoV-2 vaccination. **B-F** scRNAseq data from the reverse phenotyping dataset. The complete dataset comprising both $CD4^+$ and $CD8^+$ T cells is shown (annotation described in methods section). **B** UMAP with Leiden clusters (left panel; n=153,468 cells in total). Cluster numbers are depicted on the right with reactive cluster 27 highlighted in bold. UMAP of stimulated (blue) and unstimulated (orange) T cells (middle panel) and of $CD4^+$ and $CD8^+$ T cells (right panel). For *IFNG*, cells located within the reactive cluster are displayed with increased point size. **C** Fraction of $CD4^+$ and $CD8^+$ T cells within the reactive cluster 27. **D-E** *IFNG* expression (D) and proliferation score (E) in unstimulated (stimulated cells in grey) and stimulated (unstimulated cells in grey) T cells (left), and quantification in the stimulated condition of cells in the reactive cluster 27 versus all other clusters (right). Cells with log-normalized gene expression of 0 are shown in grey in UMAPs. Statistical testing by Mann-Whitney U test. ****p<0.0001. **F** Dot plots of log-normalized expression of representative genes per cluster. Selected genes of the reactive cluster are highlighted in grey. Numbers on the left indicate cluster number. Reactive cluster 27 is highlighted in bold.

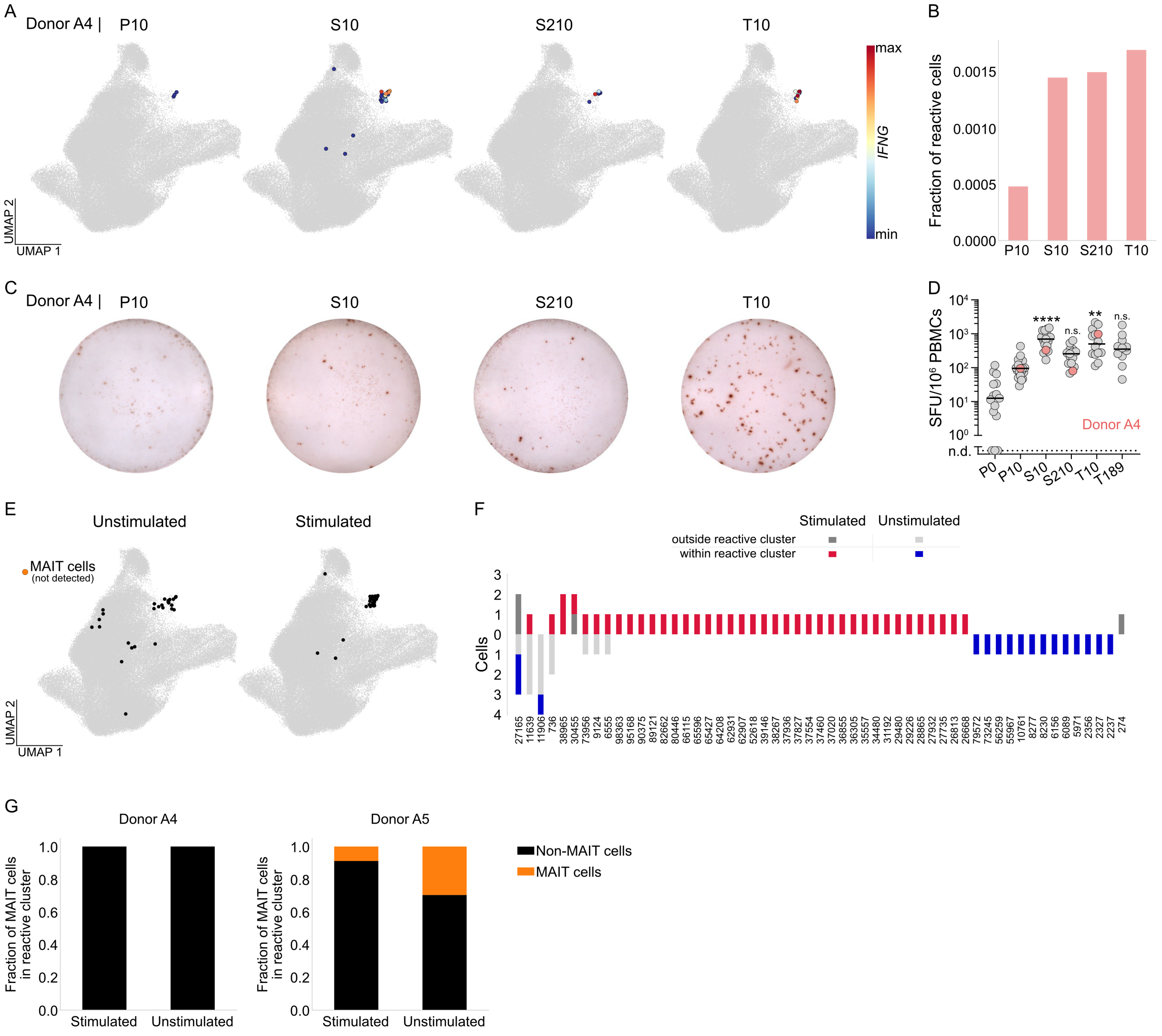


**Suppl. Fig. 2: Identification of SARS-CoV-2 spike-reactive CD4⁺ T cells in donor A4 by reverse phenotyping. A** UMAP visualization of cells classified as reactive (cells belonging to clones where at least one cell is in the reactive cluster) from donor A4 at individual time points after primary (P), secondary (S) and tertiary (T) vaccination in the stimulated condition. Color gradient indicates *IFNG* expression at indicated time points. Non-reactive cells and cells from the other donor are shown in grey. **B** Fraction of cells from donor A4 at each time point belonging to the reactive cluster. **C-D** Identification of spike-reactive T cells after 20h of *in vitro* re-stimulation of PBMCs with 15-mer peptides covering the complete wildtype spike protein. Peptides were provided in two subpools, S1 (depicted in Suppl Fig. 2C) and S2. Primary data (C) of donor A4 is shown. Quantification (D) of spot-forming units (SFU) for IFNγ ELISpot (combined frequencies of S1 and S2 subpools), data points represent individual donors (n=12-19 per time point), solid lines indicate the mean. Samples without SFU above the negative control were set to not detected (n.d.). Donor A4 is highlighted in pink. Statistical testing by Kruskal-Wallis test followed by Dunn's multiple comparisons test. Significant differences to P10 time-point are indicated. *p<0.05, **p<0.01, ***p<0.001, ****p<0.0001, n.s. not significant. **E** UMAP showing all cells classified as reactive (cells located in the reactive cluster or belonging to clones where at least one cell is in the reactive cluster) from donor A4 across all pooled time points in the unstimulated (left; n=28 cells) and stimulated (right; n=44 cells) condition. Cells classified as MAIT cells (annotation described in methods section) are depicted in orange (not detected for donor A4). Non-reactive and cells from the other donor are shown in grey. **F** Reactive clones of donor A4 are shown with the respective number of cells located within or outside the reactive cluster in the stimulated and unstimulated conditions. **G** Fraction of MAIT cells located within the reactive cluster under stimulated and unstimulated conditions for donor A4 (left) and donor A5 (right).

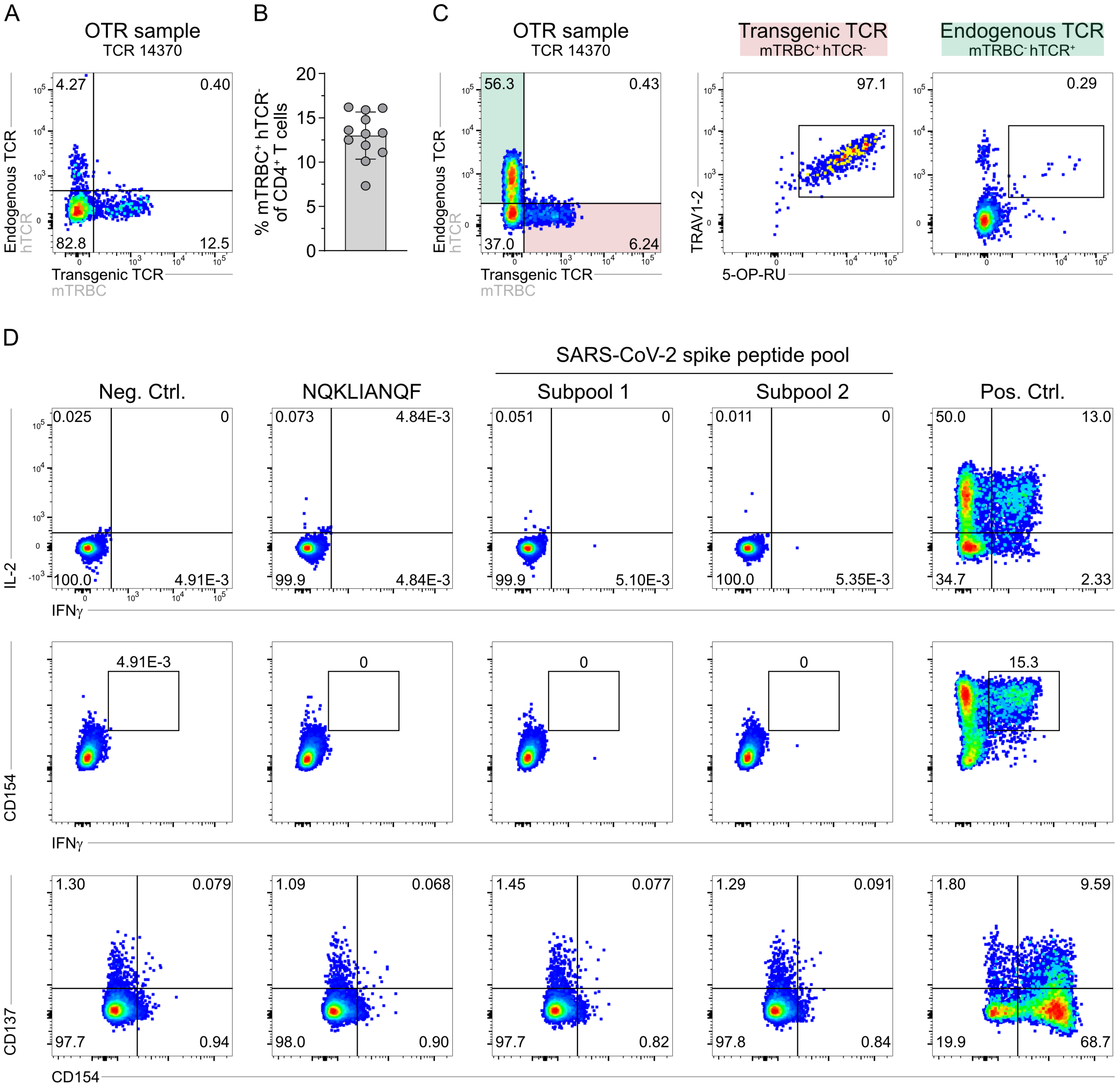


**Suppl. Fig. 3: Functional validation of MAIT TCR clone 14370 identified by scRNAseq. A-B** TCR14370 identified in donor A5, was re-expressed in primary human T cells via CRISPR/Cas9-mediated orthotopic TCR replacement (OTR) with a murine constant region (mTRBC) to be distinguishable from the endogenous TCR (hTCR). Representative flow cytometry plot (A) four days after electroporation, pre-gated on living CD4$^+$ lymphocytes. Quantification (B) of knock-in (KI) efficiency (protein expression) four days after electroporation (n=12, two independent experiments). Data points represent technical replicates, bars with error bars show the mean +/- s.d.. **C** Transgenic TCR 14370 and endogenous TCRs were stained with MR1 tetramers loaded with the MAIT ligand 5-OP-RU, alongside prototypical TRAV1-2 staining, 14 days after electroporation. Cells were pre-gated on CD19$^-$ mTRBC$^+$ hTCR$^-$ CD4$^+$ T cells (transgenic TCR, highlighted in red) or CD19$^-$ mTRBC$^-$ hTCR$^+$ CD4$^+$ T cells (endogenous TCRs, highlighted in green). **D** Transgenic T cells were co-incubated with antigen-loaded PBMCs (serving as APCs) from donor A5. PBMCs were either pulsed with $10^{-4}$ M NQKLIANQF peptide or 1 µg/mL of 15-mer peptides covering the complete wildtype spike protein (provided in two separate subpools, S1 and S2). Negative control (Neg. Ctrl) = solvent, positive control (Pos. Ctrl) = PMA/ionomycin. Reactivity was assessed by flow cytometry based on activation marker expression and intracellular cytokine staining. Flow cytometry plots are pre-gated on living CD19$^-$ CD4$^+$ hTCR$^-$ lymphocytes.

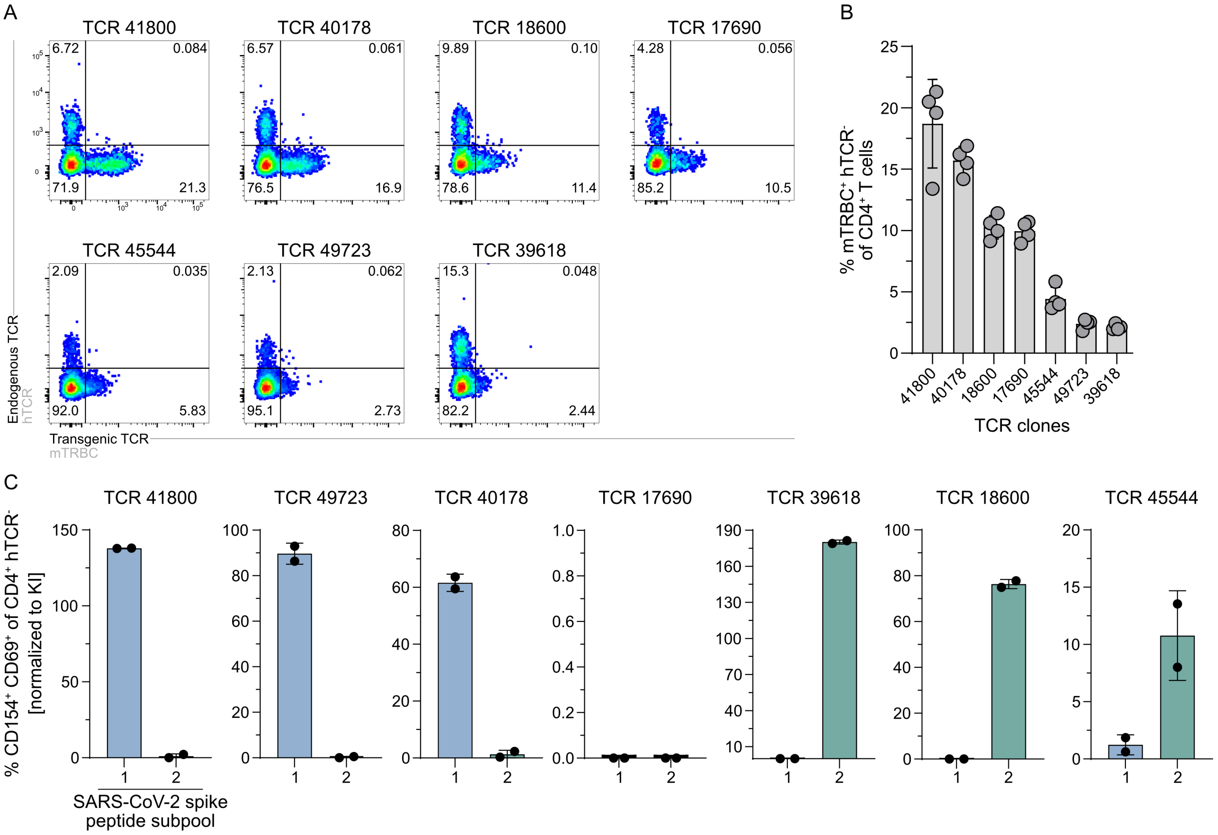


**Suppl. Fig. 4: Transgenic re-expression and functional characterization of spike-reactive TCRs identified by scRNAseq.** **A-B** Seven non-MAIT TCRs identified in donor A5 to be spike-reactive were re-expressed in primary human T cells via CRISPR/Cas9-mediated orthotopic TCR replacement (OTR) with a murine constant region (mTRBC) to be distinguishable from the endogenous TCR (hTCR). Representative flow cytometry plots (A) four days after electroporation, pre-gated on living CD4$^{+}$ lymphocytes. Quantification (B) of knock-in (KI) efficiency (protein expression) four days after electroporation (n=4, one experiment). Data points represent technical replicates, bars with error bars show the mean +/- s.d.. **C** Transgenic T cells were co-incubated with antigen-loaded PBMCs (serving as APCs) from donor A5. PBMCs were loaded with 1 µg/mL of 15-mer peptides covering the complete wildtype spike protein (provided in two separate subpools, S1 and S2) and reactivity was assessed by flow cytometry for activation marker expression. Quantification of CD69$^{+}$ CD154$^{+}$ double-positive CD19$^{-}$ CD4$^{+}$ hTCR$^{-}$ T cells per clone (normalized to mTRBC$^{+}$ cells) after SARS-CoV-2 spike-specific stimulation with S1 or S2 subpools (n=2, one experiment). Data points represent technical replicates, bars with error bars show the mean +/- s.d..

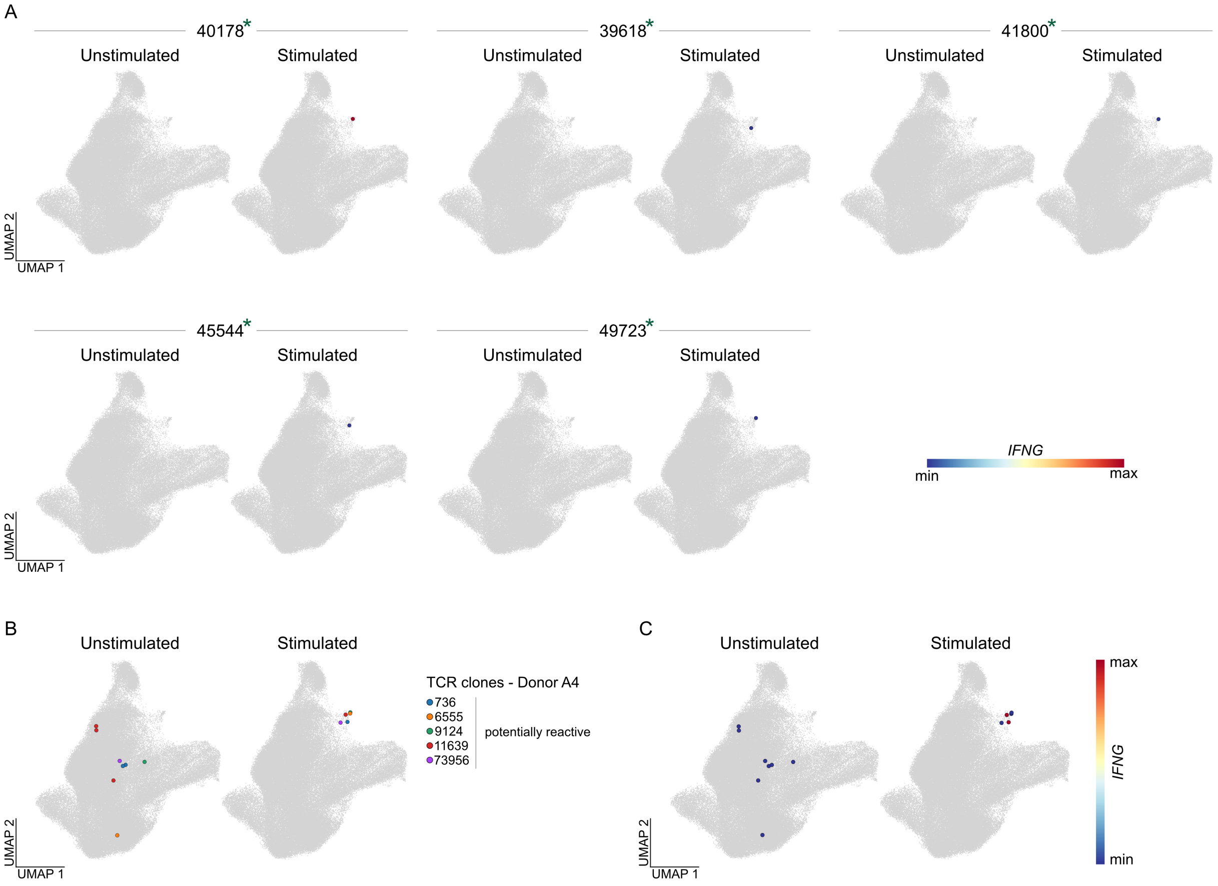


**Suppl. Fig. 5: Clonotype-specific recruitment into reactive cluster upon spike-specific stimulation. A** *IFNG* expression in unstimulated or stimulated T cells for five representative clonotypes classified as reactive in donor A5. For each clonotype, cells belonging to that clonotype are shown in an individual paired panels (unstimulated condition on the left, stimulated condition on the right), while cells not belonging to that clonotype are shown in grey. Each clone is annotated with its functionally validated SARS-CoV-2 spike reactivity status (green asterisk = reactive). For the remaining functionally tested clones, see Fig. 2E. **B-C** Based on results from Fig. 2A-E and Suppl. Fig. 2E-F, clones with cells located outside the reactive cluster in the unstimulated and within the reactive cluster in the stimulated condition were classified as potentially reactive clones. For donor A4, cells of these clones (n=5) are depicted in the unstimulated (left) and stimulated condition (right) in the UMAP (B). Colors indicate cells belonging to the same clone. *IFNG* expression in unstimulated or stimulated T cells for these clones (C). Cells not belonging to these clonotypes are shown in grey. For visualization, log-transformed expression values >3 were clipped.

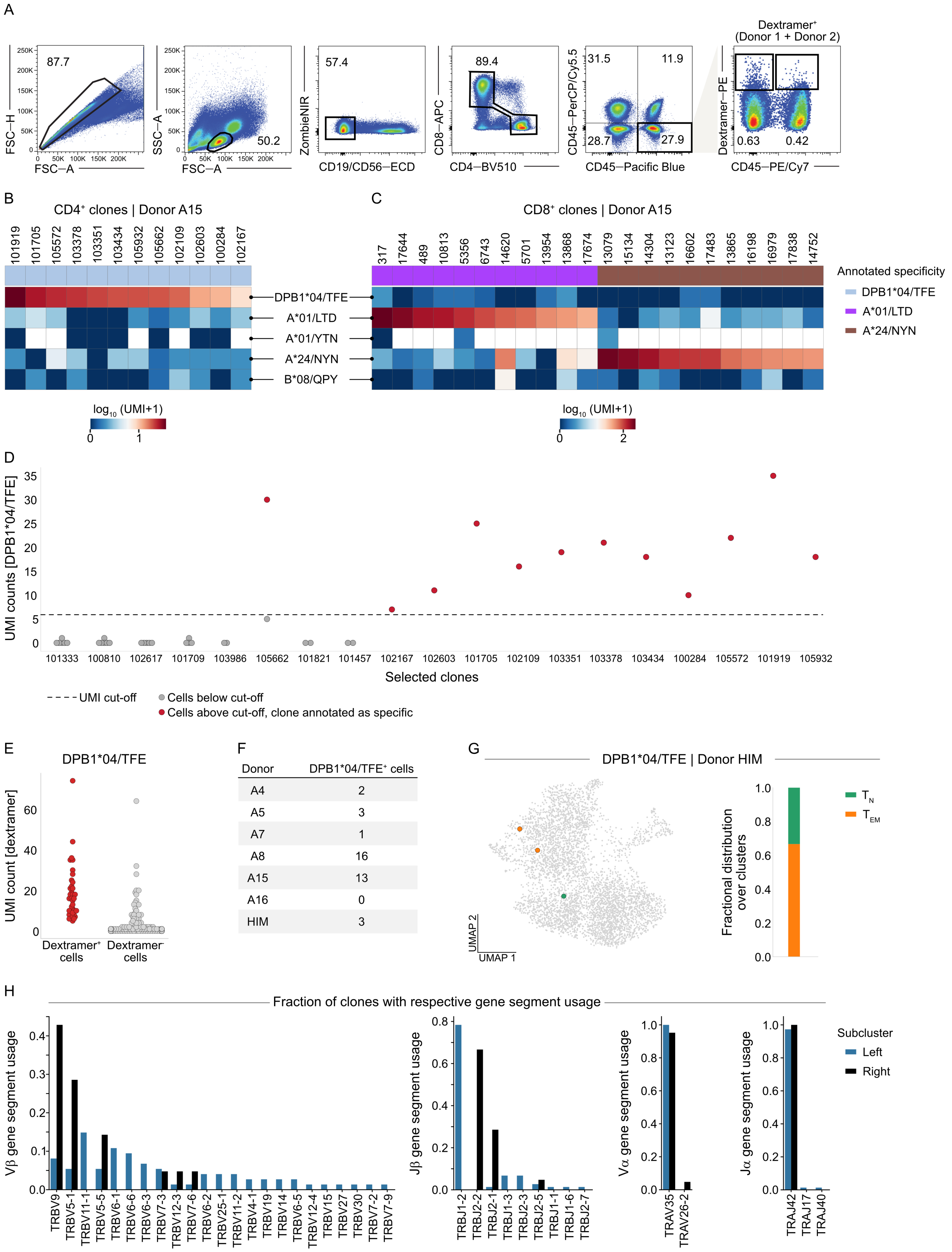


**Suppl. Fig. 6: Enrichment for DPB1*04/S$_{167}$-specific T cells using pHLA class II dextramer.**

*Legend see next page*

**Suppl. Fig. 6: Enrichment for DPB1*04/S$_{167}$-specific T cells using pHLA class II dextramer. A** Gating strategy for one representative, pooled sample to sort T cells for scRNAseq. Single, live, CD19$^-$ and CD56$^-$, CD4$^+$ or CD8$^+$, dextramer$^+$ lymphocytes were enriched for a total of seven donors across seven time points after vaccination. Based on individual CD45 color-barcodes, cells from different donors were identified in the pooled sample to ensure balanced cell numbers per donor during sorting. During later analysis, donors were identified via hashtag antibodies. The main purpose of the sort was the enrichment of dextramer$^+$ cells. **B-C** Representative heatmaps showing average UMI counts of detected clones with assigned epitope-specificity for SARS-CoV-2 spike-derived HLA class II epitope DPB1*04/S$_{167}$ (abbreviated here as DPB1*04/TFE) and HLA class I epitopes (A*01/LTD, A*01/YTN, A*24/NYN, B*08/QPY). Representative CD4$^+$ (B) and CD8$^+$ (C) T cell clones from donor A15 are shown. All epitopes are HLA-matched to donor A15. **D** Representative distributions of dextramer$^+$ and dextramer$^-$ cells from donor A15. UMI counts for the DPB1*04/TFE dextramer are shown for each cell of representative clones. The UMI cut-off was set to six and depicted as a dotted line. Individual cells with UMI counts below this cut-off are depicted in grey. Cells with higher UMI counts are shown in red if the clone was assigned dextramer$^+$ after applying additional cell purity (40%) and clone purity (50%) criteria (see Methods section). **E** Distribution of DPB1*04/TFE dextramer UMI counts of cells annotated as dextramer$^+$ and dextramer$^-$. Cells from all HLA-matched donors (n=7) are shown. **F** Number of DPB1*04/TFE dextramer$^+$ cells across all HLA-matched donors (n=7). **G** UMAP visualization (left) and quantified fractional distribution (right) of DPB1*04/TFE-specific T cells from donor HIM, 189 days after 215$^{th}$ vaccination. Colors represent cluster location of epitope-specific cells at the indicated time point. Cells without the indicated epitope-specificity are shown in grey. **H** V- and J- gene segment usage among clones located in two clusters containing our reactive and DPB1*04/TFE-specific clones together with spike-annotated published clones (left and right TCRdist clusters, see Fig. 3H). The fraction of clones with the respective gene segment usage for the TCR$\beta$ chain (two left plots) and TCR$\alpha$ chain (two right plots) is shown. For TCRdist clustering and CDR3 sequence motifs, see Fig. 3H.
